# Alleviating cell lysate-induced inhibition to enable RT-PCR from single cells in picoliter-volume double emulsion droplets

**DOI:** 10.1101/2022.08.16.504154

**Authors:** Margarita Khariton, Conor J. McClune, Kara K. Brower, Sandy Klemm, Elizabeth S. Sattely, Polly M. Fordyce, Bo Wang

## Abstract

Microfluidic droplet assays enable single-cell PCR and sequencing assays at unprecedented scale, with most methods encapsulating cells within nanoliter sized single emulsion droplets (water-in-oil). Encapsulating cells within picoliter double emulsion (DE) (water-in-oil-in-water) droplets allows sorting on commercially available FACS machines, making it possible to isolate single cells based on phenotypes of interest for downstream sequencing and PCR analyses. However, DE droplets must be within the range of large cells (<20 pl) for sorting on standard cytometers, posing challenges for molecular biology as prior reports suggest that reverse transcription (RT) and PCR amplification cannot proceed efficiently at volumes below 1 nL due to cell lysate-induced inhibition. To overcome this limitation, we used a plate-based RT-PCR assay designed to mimic reactions in picoliter droplets to systematically quantify and ameliorate the inhibition. We find that RT-PCR is blocked by lysate-induced cleavage of nucleic acid probes and primers, but that this cleavage can be efficiently alleviated through heat lysis that simultaneously inactivates inhibitory lysate components. We further show that the magnitude of RT-PCR inhibition depends strongly on cell type, but that RT-PCR can proceed in low-picoscale reaction volumes for most cell lines tested. Finally, we demonstrate one-step RT-PCR from single cells in 20 pL double emulsion droplets with fluorescence detectable via FACS. These results open up exciting new avenues for improving picoscale droplet RT-PCR reactions and expanding microfluidic droplet based single-cell analysis technologies.

## Introduction

Encapsulation of cells within microfluidic droplets enables high-throughput genomic and transcriptomic profiling of individual cells through PCR or sequencing-based techniques and has transformed many areas of biomedical research.^1-3^ In most cases, cells are encapsulated within large water-in-oil single emulsion droplets (0.5-5 nL), as cell lysis within smaller volumes is commonly thought to inhibit downstream molecular biology reactions.^4-10^ However, the ability to interrogate cells encapsulated within even smaller picoliter volume droplets would confer several key advantages. First, it would significantly reduce the probability of encapsulating multiple cells in the same droplet, thereby enhancing data quality of downstream analyses.^11,12^ Second, using picoliter droplets for single-cell analysis would dramatically lower reagent costs, as each droplet reaction requires less reagents. Finally, as cell sorting requires cells or objects be significantly smaller than the sort nozzle, picoliter volume compatibility would enable encapsulation of cells within aqueous-suspended double emulsions (water-in-oil-in-water) that can be screened and sorted using standard commercially-available fluorescence-activated cell sorter (FACS) instruments for downstream applications.^12-18^

Small droplet size, however, poses a fundamental challenge. As the size of the droplet approaches that of a large cell (<1 nL), it is widely reported that the increased concentration of cell lysate inhibits the molecular biology reactions that are the basis of all downstream analyses, including reverse transcription (RT) and PCR amplification.^4-10^ However, the precise volume at which inhibition begins to emerge has not been rigorously quantified. Although inhibition is thought to occur through cell lysate interfering with polymerases and nucleic acids (e.g., DNA or RNA templates and DNA primers) through direct binding, steric interference, premature cleavage, or sequestration of cofactors (e.g, Mg^2+^ or Mn^2+^ required for enzymatic activities)^19,20^, it remains ambiguous which step(s) of the RT-PCR reaction are inhibited and by how much. A quantitative and systematic characterization of these inhibitory effects is required to mitigate them and enable new droplet-based assays.

Here, we built on the Dropception^12^ technology to use double emulsion (DE) droplets as picoliter-volume reactors for RT-PCR from single cells with the ability to screen DE droplets based on fluorescence using commercial FACS machines. To be compatible with standard FACS instruments, droplets must pass without clogging or shearing through FACS nozzles (typically <130 µm in diameter), constraining their maximum internal volume to ∼100 pL (∼60 µm in diameter)^12,13,21^. We systematically investigated every step of RT-PCR reactions from single cells within small droplets and made several surprising findings.

First, we found that high concentrations of cellular lysate in picoscale droplets led to cleavage of DNA probes, producing a false positive signal and interfering with true amplification. We were able to mitigate this by lysing cells via heat, which simultaneously releases cell contents and inactivates enzymatic digestion. Second, we found that RT-PCR was successful in the presence of surprisingly high lysate concentrations, although the efficiency dropped for low picoliter reactions. Third, by quantifying amplification of exogenously supplemented RNA and DNA templates in the presence of lysates from a variety of human and mouse cell lines, we found that lysate-induced inhibition varied significantly between several common cell lines, exhibiting a broad range of behaviors from no inhibition of RT to strong inhibition of PCR. Together, these insights and advances allowed us to demonstrate one-step RT-PCR directly from whole cells in picoliter droplets for the first time. Our progress paves the way for developing a wide variety of new single-cell analysis technologies that can isolate and enrich target cell populations based on gene expression signatures for downstream sequencing and PCR analyses.

## Materials and Methods

### Cell culture

Jurkat cells were cultured as recommended by ATCC standards, in Roswell Park Memorial Institute (RPMI) 1640 medium (Thermo Fisher Scientific) with 10% FBS (Gibco), supplemented with 100 units/mL of penicillin, 100 µg/mL of streptomycin, and 0.25 µg/mL Amphotericin B (Gibco). HEK-293, K-562, and RAW-264 cells were cultured as recommended by ATCC standards, in DMEM with L-glutamine (ATCC) supplemented with 10% fetal bovine serum (Gibco), 100 units/mL of penicillin/streptomycin (Gibco), and non-essential amino acids (Gibco). Hep G2 cells were cultured as recommended by ATCC standards, in Eagles’ minimal essential medium (from Corning; with sodium bicarbonate, non-essential amino acids, L-glutamine and sodium pyruvate) supplemented with 10% fetal bovine serum (Gibco) and 100 units/mL of penicillin/streptomycin (Gibco). Cell viability and density measurements were taken using a Trypan blue exclusion assay.

### Device fabrication and preparation

The Dropception device design, fabrication, preparation, and operation took place as described in Ref. 12. Briefly, devices were designed in AutoCAD 2019 (AutoDesk) and molding masters were fabricated via multilayer photolithography using SU8 photoresists (Microchem) according to the manufacturer’s instructions on 4” test wafers (University Wafer). PDMS devices were cast from these models via soft lithography using RTV 615 (R.S. Hughes).

### Double emulsion generation

Double emulsion droplets were generated using four syringe pumps (PicoPump Elite, Harvard Apparatus) for cell suspension and inner, oil, and outer sheath solutions. The oil phase was composed of HFE7500 fluorinated oil (Sigma) and 2.2% Ionic PEG-Krytox (FSH, Miller-Stephenson)^22^. The carrier phase contained 1% Tween-20 (Sigma) and 2% Pluronic F68 (Kolliphor 188, Sigma) in 1×PBS. Prior to droplet loading, cells were stained with Calcein AM UltraBlue or DeepRed (AAT Bioquest), then resuspended to a concentration of 2.4×10^6^/mL in 1× PBS supplemented with 20% OptiPrep and 0.04% BSA. Unless otherwise noted, the second inner phase contained 0.5× SuperScript III One-Step RT-PCR System buffer, 60 µl/mL SuperScript III RT/Platinum Taq High Fidelity mix (Thermo Fisher Scientific), 1% BSA (Fisher Scientific), 2 U/µL RNasin Ribonuclease Inhibitor (Promega), 1.2 µM PrimeTime primers, and µM PrimeTime 5’ FAM/Zen/3’ IBFQ probe (IDT). Each phase was loaded into syringes (PlastiPak, BD) as described in Ref. 12 and connected to the device via PE/2 tubing (Scientific Commodities). Typical flow rates were 400:100:105:6000 µL/hr (oil:cells:reagents:outer).

### Double emulsion flow cytometry

DE droplets were analyzed via FACS as described in the prior sdDE-FACS^13^ and Dropception^12^ studies. 100 µL of droplets were resuspended in 500 µL of sheath buffer containing 1% Tween-20 (Sigma) in 1×PBS and analyzed on the SH800 flow cytometer (Sony) using a standard 408 nm laser configuration with a 130 µm nozzle. After a brief pickup delay time of ∼2 min, DEs were gated on FSC-H vs. FSC-W and analyzed for Calcein AM and TaqMan FAM signal. All flow parameters are as previously reported^14^. FACS data were downsampled to 40,000 DEs and analysis performed with FlowJo software (v10).

### Droplet and plate assay RT-PCR

DE droplets were thermal cycled in individual PCR tubes. Given the shell permeability of DE droplets, we resuspended a 10 µL droplet pellet in 25 µL 2× SuperScript III One-Step RT-PCR System buffer (unless otherwise noted) to transport key salt components in the buffer across the shell. RT-PCR reactions were performed at the following conditions: 60°C 90 min RT followed by 0 to 50 cycles of 94°C 15 sec, 50°C 30 sec, 68°C 30 sec. We noticed that droplet packing appeared to alter efficiency, with smaller numbers of droplets showing enhanced activity even when the ratio of droplet volume to total volume was kept constant (**Fig. S1**); therefore, all droplet reactions were performed using total droplet pellet volumes of 10 µL.

Plate RT-PCR assays were performed with the same temperature conditions on a CFX96 Real-Time PCR Detection System (BioRad) or an Applied Biosystems QuantStudio 3 (ThermoFisher) system with FAM fluorescence readout. Unless otherwise noted, the reagent mixture was composed of 1× SuperScript III One-Step RT-PCR System buffer, 30 µl/mL SuperScript III RT/Platinum Taq High Fidelity mix (Thermo Fisher Scientific), 0.5% BSA (Fisher Scientific), 1 U/µL RNasin Ribonuclease Inhibitor (Promega), 0.6 µM PrimeTime primers, 0.3 µM PrimeTime 5’ FAM/Zen/3’ IBFQ probe (IDT), and concentrated cell suspension containing the indicated number of cells, for a final reaction volume of 10 µL. The concentrations of RT and Taq enzymes were higher than those used in standard RT-PCR reactions, as we noticed that higher concentrations improved the efficiency under conditions of high cell lysate concentrations. Cell suspensions were counted manually then diluted to reach the appropriate concentration in 1× PBS supplemented with 0.04% BSA. mTurqouise DNA was synthesized as a gBlock (IDT) and amplified by PCR. mRNA was generated by *in vitro* T7 transcription (HiScribe™ T7 ARCA mRNA Kit, NEB). DNA and mRNA at the indicated concentrations were added to the mixture of cells and RT-PCR reagents immediately before proceeding to heat lysis.

Primer sequences used in this study were as follows: human *GAPDH* forward primer, TGTAGTTGAGGTCAATGAAGGG; human *GAPDH* reverse primer: ACATCGCTCAGACACCATG; human *GAPDH* probe: /56-FAM/TTCACTGCA/ZEN/TTGCCGGACAAATCG/3IABkFQ; s*med-pc2* (prohormone convertase-2^23^ of the planarian *Schmidtea mediterranea*) forward primer: GGTGTCGGAACTGTGGATTTA; s*med-pc2* reverse primer: GCTTCAAGAGCAAGAGCAAAC; s*med-pc2* probe: /56-FAM/ACATCATTC/ZEN/GGGAACTTCTGCTGCT/3IABkFQ; mTurquoise forward primer, CGCGGACCATTACCAACAAAATACG; mTurquoise reverse primer, AACTCATCCATTCCGAGCGTGAT; mTurquoise probe, /56-FAM/TCTTGCTGG/ZEN/AGTTTGTTACCGCCGCTGGCA/3BHQ_1/. *GAPDH* primers were designed to span a 1,632 bp intron in order to prevent amplification from genomic DNA.

### Image acquisition and analysis

To extract a scalar value (C_i_, or cycle of inflection) from RT-qPCR and qPCR data, raw curves were fit to an asymmetric sigmoid (scipy: optimize package)^24^:

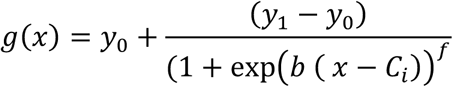

The asymmetry parameter (*f*) was fixed at 0.3, which is the approximate value for most fits without fixing *f*, to improve the stability of curve fitting.

All images were captured on a Zeiss Axio Observer Z1 inverted microscope equipped with a Zeiss Axiocam 503M camera using a 5x/0.25 objective and analyzed using the Zen Blue software (Zeiss).

## Results and Discussion

### The Dropception single-cell RT-PCR workflow

Our workflow (**Fig. 1**): (1) co-encapsulates cells, RT-PCR reagents, and transcript-specific Taqman probes within 20 pL DE droplets, (2) lyses cells, reverse transcribes mRNA, and detects transcripts of interest via Taqman PCR, and then (3) identifies and selects for droplets containing cells and transcripts based on the Taqman fluorescence signal via FACS. During encapsulation, we co-loaded cells stained with a cell viability dye (Calcein AM) and suspended within a density gradient medium to improve loading consistency (20% OptiPrep) with a solution containing One-Step SuperScript III/TaqMan RT-PCR reaction mixture and a transcript-specific primer pair and Taqman probe, and bovine serum albumin (BSA) (for enzyme and droplet stabilization). After encapsulation and cell lysis, we performed reverse transcription followed immediately by PCR via High Fidelity Platinum Taq. Amplification of transcripts of interest should lead to Taqman probe cleavage and an increase in fluorescence within the droplet, making it possible to isolate these droplets via FACS.^12,13^ After isolation, individual droplets can be air lysed into well plates; for large droplet populations (≥10,000 droplets), droplets can be demulsified using 1H,1H,2H,2H-perfluoro-1-octanol (PFO).^12,13^ This approach should allow for researchers to select cells that express a transcript of interest for downstream assays such as RNA sequencing.

**Figure 1:**
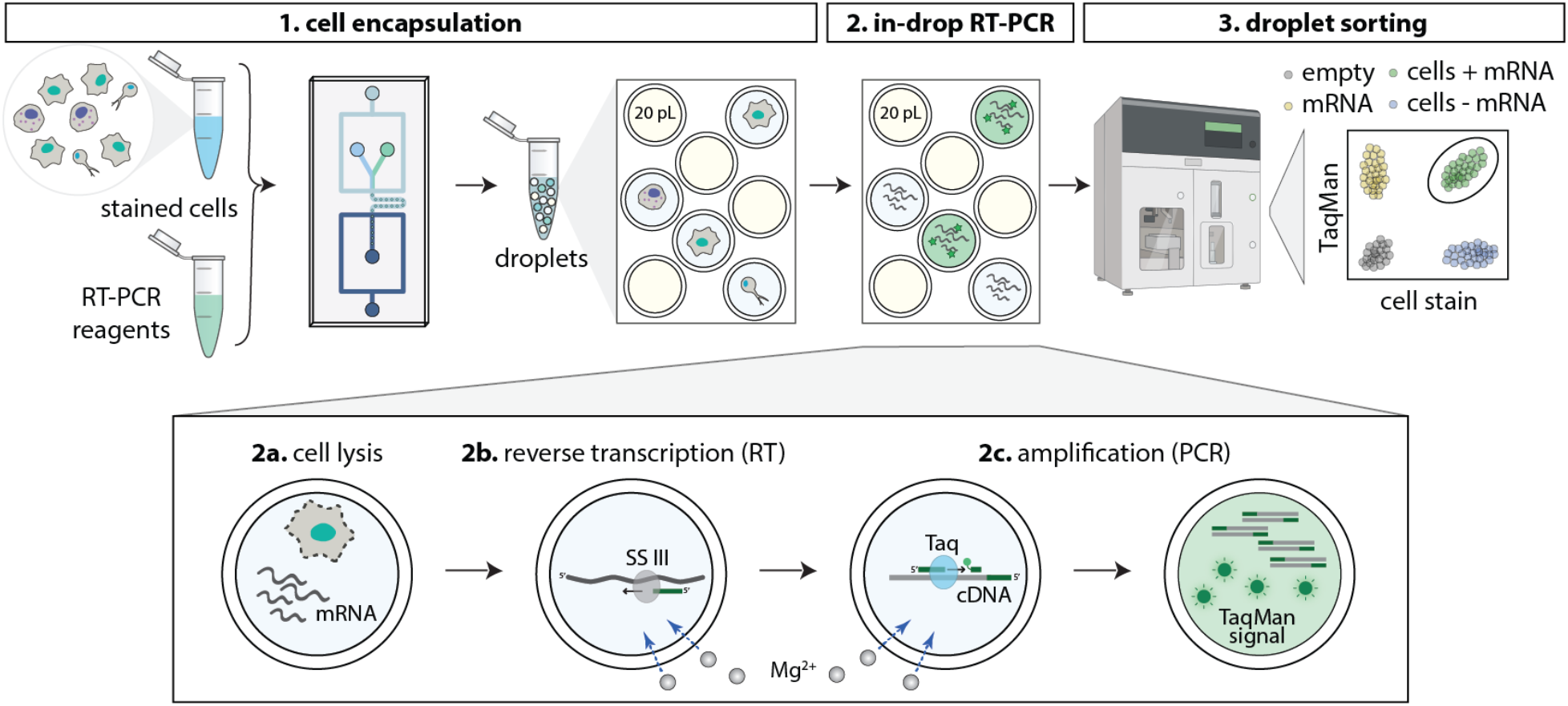
Workflow for single-cell RT-PCR in picoliter DE droplets. The workflow takes place via three main stages: (1) Calcein-stained cells are co-loaded into a single-layer microfluidic device with a TaqMan RT-PCR reagent mixture to form double emulsions (DEs) of 20 pL internal volume surrounded by an aqueous carrier solution; (2) DEs are collected, pooled, and heated to drive RT for the targeted gene using SuperScript III RT enzyme followed by 20-50 cycles of PCR by Taq DNA polymerase; (3) DEs can be analyzed and sorted via standard flow cytometry to identify Calcein-positive, TaqMan-positive droplets containing cells expressing a transcript of interest.

We anticipated four distinct categories of droplets: (1) non-fluorescent empty droplets containing neither cells nor gene target, (2) Calcein AM-positive, TaqMan-negative droplets containing cells that do not express the targeted gene, (3) Calcein AM-negative, TaqMan-positive droplets in which stray mRNA transcripts present in solution are amplified in the absence of a cell, and (4) double-positive droplets containing a cell expressing the targeted gene.

To establish that this pipeline is compatible with standard RT-PCR as previously established,^25-30^ we first tested the ability to amplify a housekeeping gene *GAPDH* from purified human RNA encapsulated within droplets (1 pg/droplet). While DE droplets were stable after thermal cycling, encapsulation of RNA with buffers typically used for RT-PCR within single emulsion droplets failed to generate fluorescence even after 50 cycles of amplification (**Fig. 2a**). As RT and PCR enzymes are critically sensitive to the concentrations of ion cofactors (e.g., MgCl_2_, MgSO_4_, and Mn) and these ions can cross thin DE oil shells,^18,31^ we reasoned that successful RT-PCR in DE droplets may require increasing ionic concentrations in the external aqueous carrier phase. Consistent with this, supplementing the external PBS carrier solution with MgCl_2_ or MgSO_4_ dramatically enhanced reaction efficiencies to ∼85% and >95%, respectively (**Figs. 2b-d, S1**). As both Taq and SuperScript III can utilize MgCl_2_ but MgSO_4_ is a more efficient cofactor for Taq,^32^ this enhancement likely results from increased Taq efficiency.

**Figure 2:**
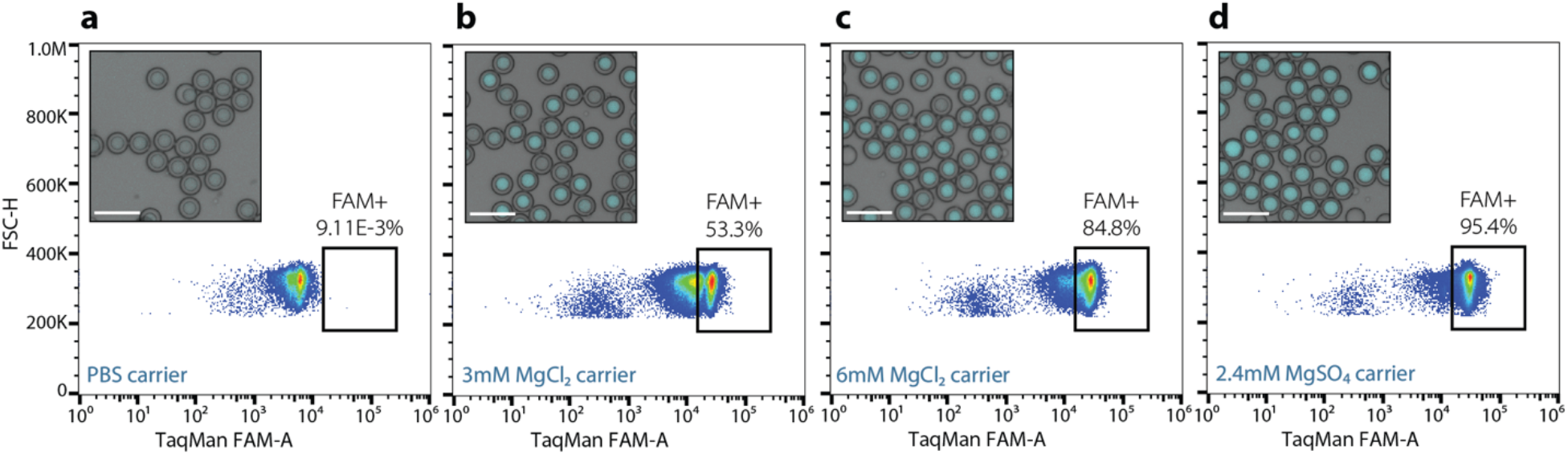
The efficiency of RT-PCR from purified RNA in DEs depends on the salt concentration of the carrier solution. FACS plots of DEs loaded with 1 pg/droplet K-562 cell total RNA and reagents for *GAPDH* RT-PCR after 50 PCR cycles when incubated in: **(a)** 1x PBS with 2% Pluronic F68 and 1% Tween-20, **(b)** 1 M-MLV RT buffer (containing 3 mM MgCl_2_), **(c)** 1 M-MLV RT buffer supplemented with an additional 3 mM MgCl_2_, and **(d)** 2 Invitrogen One-Step RT-PCR buffer (containing 2.4 mM MgSO_4_). The FAM^+^ gate is shown with the fraction of FAM^+^ droplets denoted; insets show representative merged bright field and FAM fluorescence images of droplets for each sample. Scale bars: 100 μm.

### Picoliter-volume droplet RT-PCR reveals lysate-induced DNA probe cleavage

Based on many prior studies reporting significant inhibition of RT-PCR within small (<1-5 nL) reaction volumes,^4-10^ we expected cell lysis and subsequent standard RT-PCR within 20 pL volume DEs to yield no signal. Unexpectedly, DEs loaded with human Jurkat T cells and TaqMan RT-PCR reagents to detect *GAPDH* displayed fluorescence in ∼80% of cell-containing droplets after only 20 cycles of PCR (**Fig. 3a**). However, droplets loaded with human Jurkat T cells and a probe specific to a gene that does not exist in the loaded cell (a planarian flatworm gene^21^ not present in the human genome) also generated a strong fluorescence signal (**Fig. 3b**), suggesting that the signal was not produced through PCR amplification of target genes. Indeed, we detected similar fluorescence in cell-containing droplets even without thermal cycling (**Fig. S3a**) and in the absence of both Taq and SuperScript III enzymes (**Fig. S3b**). This signal disappeared in the absence of cells (**Fig. 3c**) or TaqMan probes (**Fig. 3d**), indicating that the fluorescence readout results from an interaction between cell lysate and TaqMan probes.

**Figure 3:**
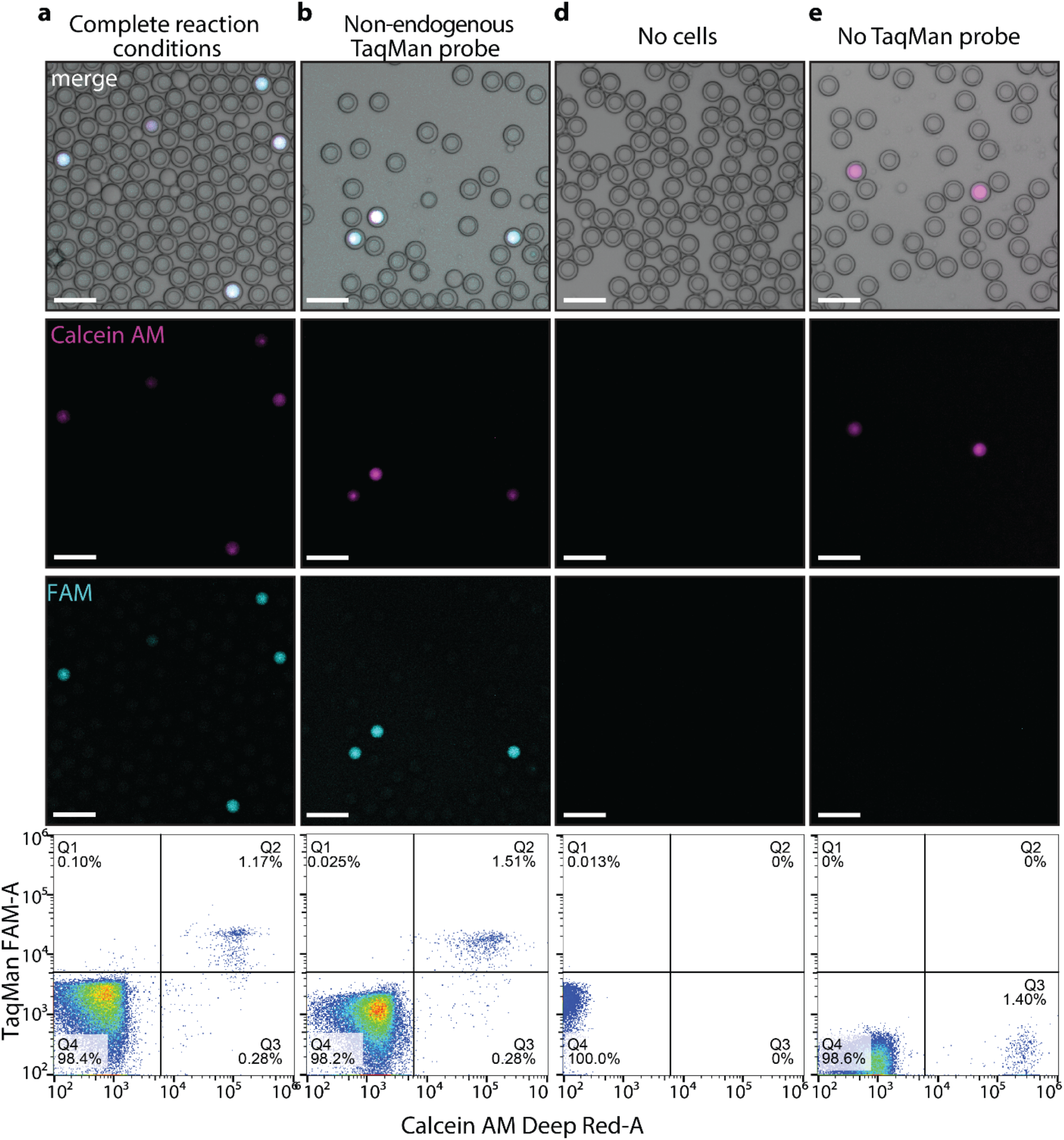
Lysate-induced probe cleavage within DE droplets. Representative bright field and fluorescence microscopy images (top; Calcein stain in magenta and TaqMan probe signal in cyan) and FACS analysis (bottom) for DE droplets loaded with: **(a)** Calcein AM Deep Red-stained Jurkat T cells and complete reaction buffer (*GAPDH* TaqMan RT-PCR reagents and lytic detergents: 0.1% NP-40, 0.1% Tween-20, 0.01% Digitonin) after RT and 20 cycles of PCR, **(b)** Calcein-stained Jurkat T cells and complete reaction buffer containing a TaqMan probe and primers for a gene absent within Jurkat cells (*Smed-pc2*, cyan) after RT and 20 cycles of PCR, **(c)** complete reaction buffer alone without cells, **(d)** Calcein-stained Jurkat T cells and reaction buffer without TaqMan probe and primers. FACS density plots display DE-gated droplets: Q1, FAM^+^ Calcein^−^; Q2, FAM^+^ Calcein^+^; Q3, FAM^−^ Calcein^+^; Q4, FAM^−^ Calcein^−^ DEs. DE gates for all FACS data are shown in **Fig. S2**. Scale bars: 100 µm.

Unlike commonly expected mechanisms of PCR inhibition, which present as diminished or absent signal, this false positive signal is consistent with non-specific cleavage of the probe and release of the fluorophore from its quencher. Supporting this notion, the cell lysate-induced probe signal also depended on the concentration of ions within the external aqueous phase. Droplets suspended in Mg^2+^-free buffers yielded weak signals (**Fig. S3c**) implicating probe cleavage via endogenous cellular enzymes (e.g., DNases).

### Heat inactivation alleviates premature probe cleavage

To enable more rapid and systematic testing and troubleshooting of small-volume conditions, we next tested whether the lysate-induced probe cleavage observed in droplets could be replicated within multiwell plates containing high effective cell lysate concentrations^8^ (**Fig. 4a**). To mimic the volume ratio within DEs, we generated 10 µL reactions containing 500,000 cells, yielding an effective lysate concentration equivalent to a single lysed cell in a 20 pL droplet. The inclusion of even gentle nonionic lysis detergents such as NP-40 (Nonidet P40)^33^ led to instantaneous probe cleavage even in the absence of amplification, observable as a high baseline fluorescence prior to thermal cycling, consistent with lysate-mediated digestion of DNA primers (**Fig. 4b**). In contrast, cells incubated on ice with RT-PCR reagents in the absence of NP-40 showed no probe cleavage until the start of the amplification reaction, suggesting that replacing detergent-mediated lysis steps with alternative methods could improve RT-PCR results by eliminating premature probe cleavage (**Fig. 4b**).

**Figure 4:**
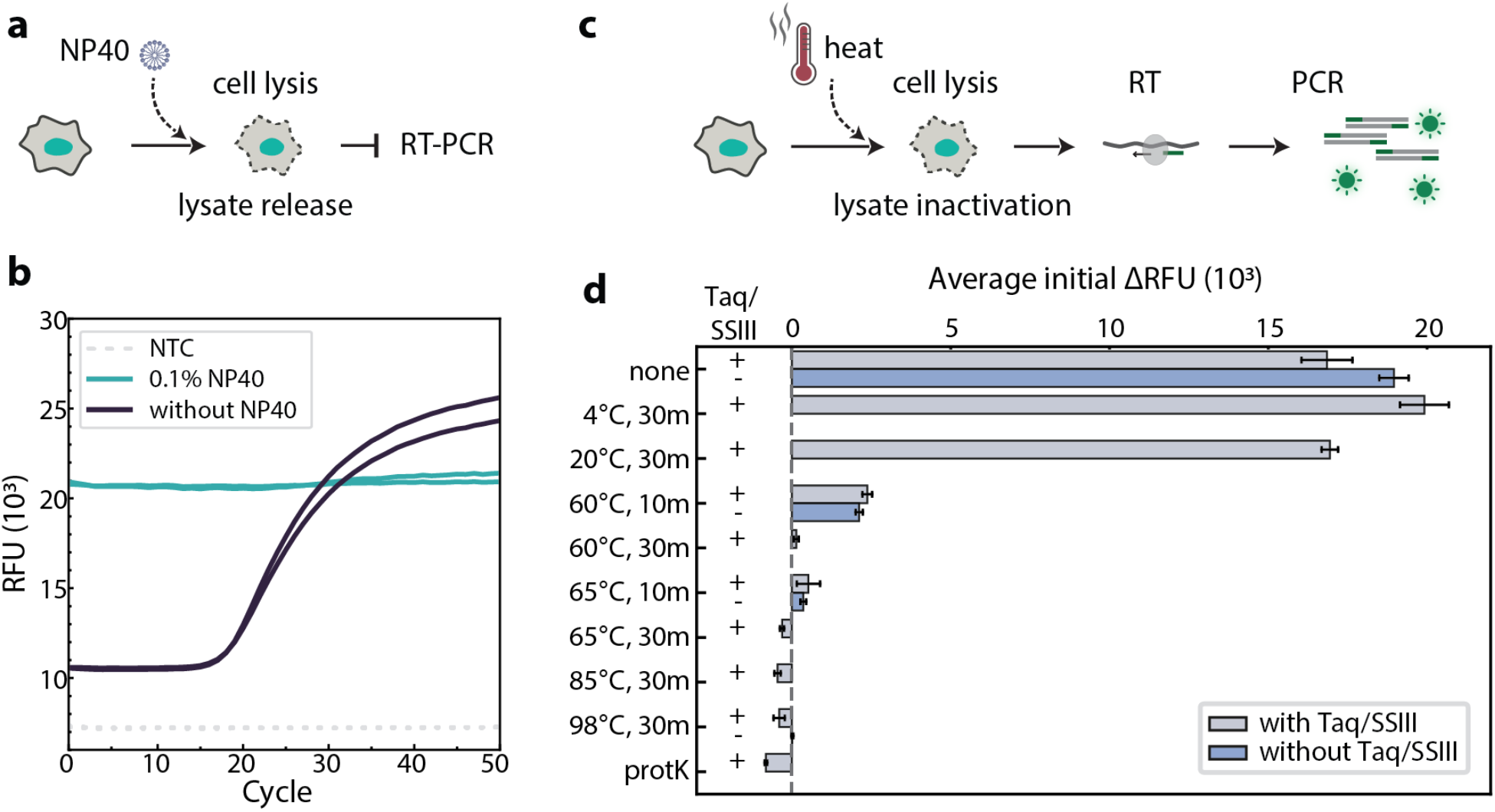
Heat inactivation mitigates lysate-induced probe cleavage. **(a)** Schematic illustrating cell lysis and subsequent lysate release upon addition of NP-40 detergent, inhibiting RT-PCR. **(b)** qPCR traces of plate-based TaqMan *GAPDH* RT-PCR reactions at high effective cell concentrations (500,000 Jurkat T cells per 10 µL reaction, equivalent to 1 cell in a 20 pL DE). Reactions containing 0.1% NP-40 exhibit high initial relative fluorescence unit (RFU) prior to PCR, indicating premature probe cleavage, and no amplification. NTC: no template control. **(c)** Schematic illustrating an initial heat treatment of cells prior to RT-PCR to simultaneously lyse cells and inactivate cleavage inducing factors in cell lysate. **(d)** Average initial RFU values of the first three PCR cycle measurements (normalized to no probe control) with detergent-free conditions for 500,000 cells per 10 µL reaction; probe cleavage is visible as a high initial RFU value. Cells were incubated at the indicated temperatures and times or with 0.5 mg/mL proteinase K (65°C, 15 min incubation followed by 85°C, 15 min inactivation) prior to combining with the reaction mixture. The color of bars indicates whether reactions took place in the presence (+, grey) or absence (–, blue) of Taq and SuperScript III enzymes; error bars indicate standard deviation.

Prior reports suggested that heat treatment alone can lyse cells.^33^ We therefore attempted to identify conditions capable of driving efficient lysis and simultaneously inactivating endogenous DNases without denaturing the other enzymes required for RT and PCR (**Fig. 4c**). While incubation at either 4 °C or 20 °C for 30 min had no effect on premature probe cleavage, which is visible as high fluorescence intensity prior to amplification, incubation at 60°C or higher for 30 min or longer largely eliminated it (**Fig. 4d**), consistent with the fact that non-thermophilic endogenous proteins typically denature between 40-70 °C.^34^ We therefore chose to heat inactivate at 60 °C for 90 min to ensure maximal cell lysis prior to amplification for all subsequent assays; importantly, these conditions are also optimal for reverse transcriptase activity, permitting a single step for continuous lysis and cDNA production. However, after 60°C for 90 min, we still observed a slightly decreased dynamic range for the fluorescence signal due to small amounts of lysate-induced probe cleave prior to heat inactivation.

### Distinct regimes of whole-cell RT-PCR at a range of reaction scales

Our plate-based assay allowed us to efficiently vary cell lysate concentration and quantify RT-PCR efficiency (**Fig. 5a**). To quantify amplification from qPCR curves with differing baseline and saturation values in an unbiased manner, we fit curves to an asymmetric sigmoid and used the cycle of inflection point (C_i_), instead of C_t_, cycle threshold, which was difficult to consistently quantify across curves with different shapes and amplitudes as previously noted^24^. With Jurkat T cells, we found that C_i_ scaled non-linearly with increasing cell numbers over several regimes (**Fig. 5b**). Reactions containing 30,000 cells or fewer in 10 µL (the lysate concentration equivalent of a single cell in a droplet larger than 300 pL) followed the expected log-linear trends in C_i_ in parallel to those of purified RNA samples. Below the equivalent droplet size of 300 pL, C_i_ values began to plateau. At even higher concentrations (>328,000 cells in 10 µL, equivalent to a cell in a 30 pL droplet), C_i_ increased with increasing cell numbers, suggesting that within this ‘inhibition’ regime, lysate-induced inhibition began to dominate and block the reaction. However, even at these very high lysate concentrations, amplification traces and C_i_ values indicate successful reactions, contrary to extensive prior claims of pervasive inhibition in small volumes.^4-10^

**Figure 5:**
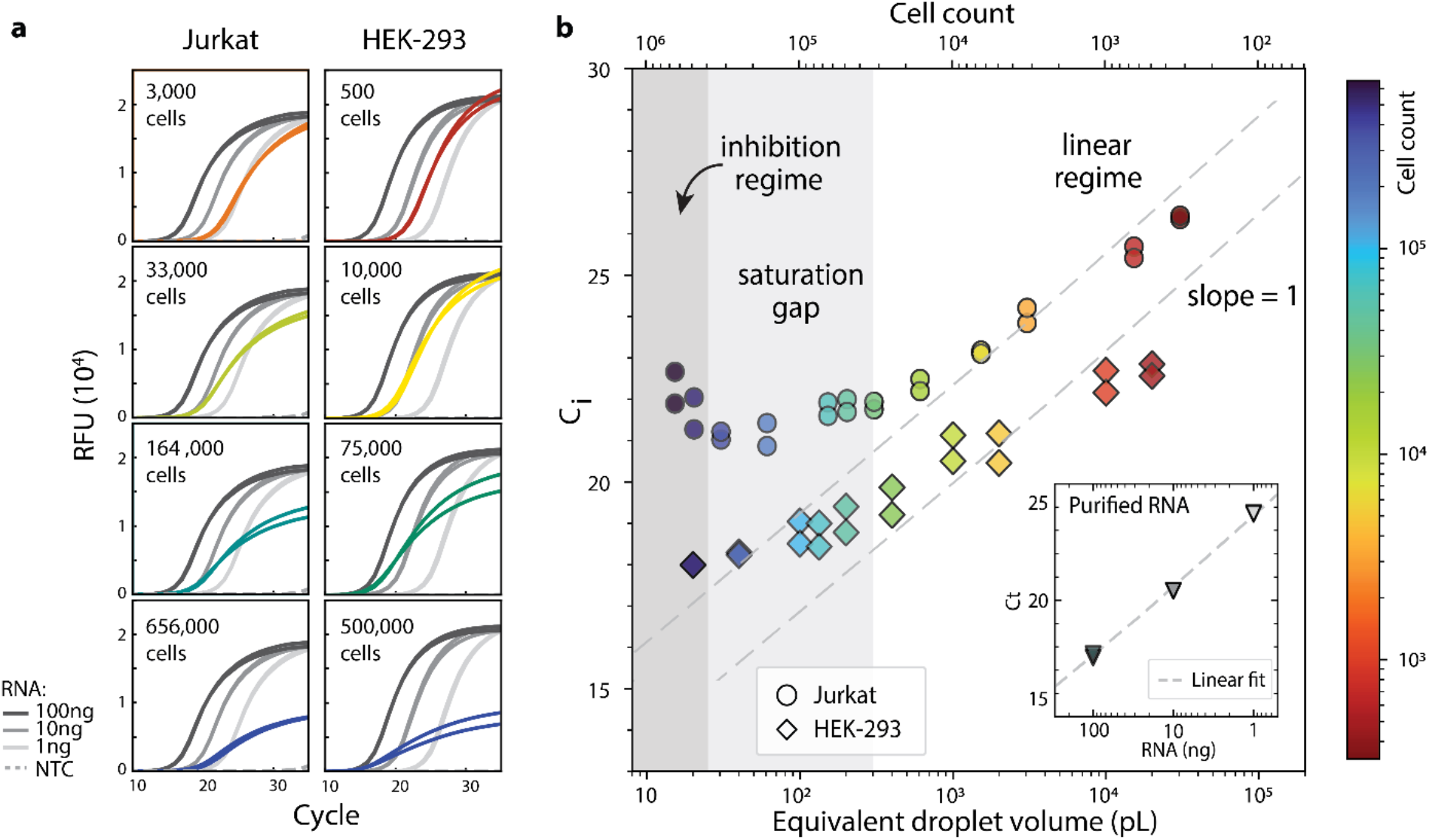
RT-PCR reaction efficiency depends on cell lysate concentration. **(a)** Baseline-subtracted qPCR traces for 10 µL reactions containing between 500 cells (equivalent to 1 cell in a 400 nL droplet) and 656,000 cells (equivalent to 1 cell in a 15 pL droplet) detecting *GAPDH* transcripts (colored traces). Left: Jurkat T cells, right: HEK-293 cells. Cell were lysed via heat. Gray lines: amplification curves for purified Jurkat RNA dilution controls; dashed lines: no template control (NTC) with neither cell nor RNA added. **(b)** C_i_ values for a range of cell counts in a 10 µL reaction plotted as a function of the equivalent single cell-loaded droplet volume. C_i_ values for Jurkat T cells (circles) decrease loglinearly with increasing cell counts until 30k cells (linear regime), then saturate until 328,000 cells (saturation gap, light gray shading), after which reaction inhibition increases C_i_ (inhibition regime, dark gray shading). In contrast, HEK-293 cells (diamonds) exhibit much lower inhibition, maintaining log-linear behavior until 100 pL equivalent volume and never entering an inhibition regime. Inset: Jurkat purified RNA dilution controls. Dashed lines: theoretical log-linear relationship with C_i_ and cell count with a slope of 1.

By contrast, human embryonic kidney (HEK-293) cells only showed a slight deviation off the log-linear trend when cell number exceeded 100,000 (equivalent to a cell in a 100 pL droplet), indicating moderate RT-PCR inhibition, and even higher effective lysate concentrations did not yield an inhibition regime in which Ci values increases with increasing cell concentrations (**Fig. 5a,b**). These results establish that RT-PCR directly from a single cell in picoscale volumes is possible and suggest that the lysate-induced inhibition is not ubiquitous and significant but rather depends on cell type.

### Quantifying the degree of inhibition of RT-PCR

Understanding the source of inhibition is critical for identifying strategies to alleviate it and for assessing feasibility of different downstream applications. For example, efficient cell lysis and RT is already sufficient for constructing RNA-seq libraries in low-volume droplets, while sorting and sequencing only droplets containing cells expressing a target transcript requires efficient RT and in-drop PCR. We therefore attempted to quantify the relative contributions of each step of RT-PCR to its reduced overall efficiency.

Since heat is reported to be an inefficient means of cell lysis,^33^ we first tested if RT efficiency is limited by incomplete lysis, leading to inaccessible endogenous RNA. For this, we pre-incubated 500,000 Jurkat T cells at 60°C to disable lysate-induced probe cleavage, added additional NP-40 detergent prior to combining with RT-PCR mixture, and quantified the initial fluorescence signal and RT-PCR efficiency (**Fig. S4a,b**). The addition of 0.1% NP-40 did not improve amplification even though this concentration of NP-40 is documented to be highly effective across single- and bulk-cell sequencing and RT-PCR assays.^8,26,33^ Furthermore, mechanical shearing of cells by pipetting did not improve amplification (**Fig. S4c,d**). These data suggest that cell lysis was not the limiting factor of our assay. To ensure that the amplification efficiency loss was not simply due to RNA hydrolysis during the heat pre-treatment, we amplified purified RNA via RT-PCR with and without a 60°C pre-treatment step and confirmed that efficiency was unchanged by pre-treatment (**Fig. S4e,f**).

Next, we examined the efficiency of the RT and PCR steps separately. To determine the degree to which PCR alone is inhibited in the presence of cell lysate, we spiked known amounts of an exogenous DNA (*mTurquoise)* not present within Jurkat T cells into 10 µL reactions containing up to 500,000 cells (**Fig. 6a**) and quantified amplification using C_i_ (**Fig. S5**).

**Figure 6:**
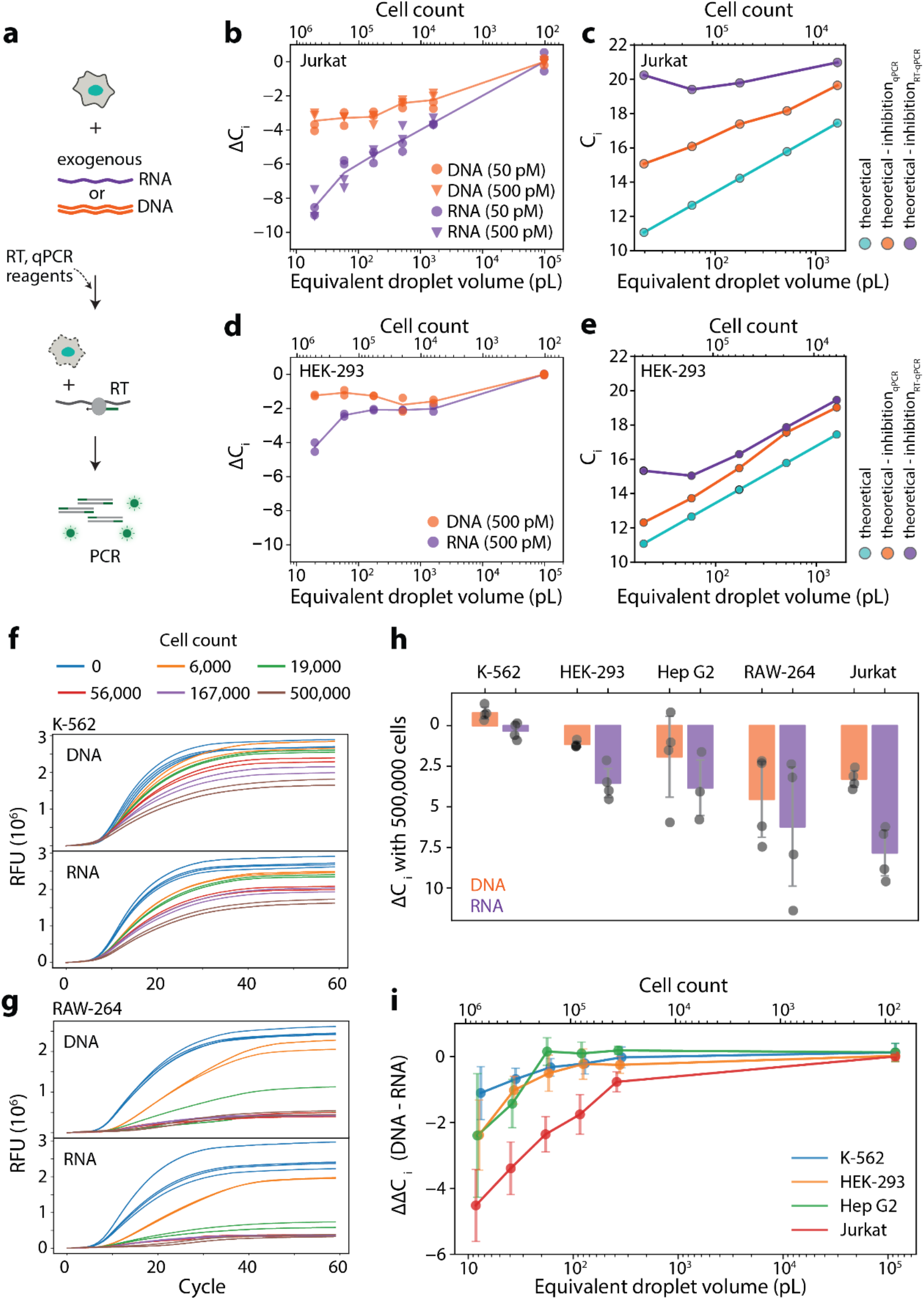
Quantifying the degree to which RT-PCR is inhibited at RT and PCR steps using exogenously supplied DNA and mRNA. **(a)** Schematic of assay quantifying lysate-induced inhibition of PCR or RT-PCR using exogenous supplemented DNA and mRNA respectively. **(b)** Inhibition of PCR (orange) or RT-PCR (purple), quantified as changes in *C*_i_ (ΔCi) for exogenous DNA and mRNA templates in the presence of Jurkat T cells designed to mimic small volume reactions. **(c)** Theoretically expected C_i_ for endogenous transcripts whose abundance scales linearly with cell number (cyan); subtracting expected inhibitory effects on PCR (orange) and on RT-PCR (purple) measured in (**b**), which reproduces observed amplification regimes (inhibition, saturation and linear regimes) for Jurkat T cells shown in **Fig. 5b. (d)** Measured inhibition of PCR and RT-PCR vs. HEK-293 cell concentration. **(e)** Expected C_i_ of endogenous transcripts for HEK cells, showing no inhibition regime. **(f-g)** Baseline-subtracted qPCR traces with exogenous DNA or mRNA templates with increasing concentrations of K-562 (**f**) and RAW-264 (**g**) cells. **(h)** ΔCi between 0 and 500,000 cells for five cell lines quantified using either exogenous DNA (PCR) or mRNA (RT-PCR) templates. **(i)** ΔΔC_i_, the difference between inhibition measured using DNA vs. mRNA templates, quantifies inhibition to RT as a function of cell concentrations for four cell lines. RAW-264 cells were excluded due to complete inhibition of the PCR reaction at high lysate concentrations.

Regardless of the template concentration, qPCR signal amplification lagged by ∼2 cycles with 6,000 cells in 10 µL (a lysate concentration equivalent to a single cell in a 2 nL droplet) and ∼4 cycles with 500,000 cells (equivalent to a single cell in a 20 pL droplet) (**Fig. 6b**). In parallel, we spiked in known amounts of *mTurquoise* mRNA and found that the RT step was inhibited to a degree comparable to that of PCR: amplification was further delayed by ∼2 cycles with 6,000 added cells and ∼5 cycles with 500,000 cells (**Fig. 6b**). These data explain the linear, saturation, and inhibition regimes observed when amplifying endogenous transcripts from Jurkat T cells. Without inhibition, C_i_ for RT-PCR of an endogenous transcript should increase loglinearly with the number of cells. Subtracting the measured loss from the expected C_i_ (**Fig. 6c**) reproduced the observed relationship between cell concentration and amplification with *GAPDH* probes (**Fig. 5b**).

In contrast, HEK-293 cell lysate was much less inhibitory, even though they have a ∼50% larger volume compared to Jurkat T cells implicating higher lysate concentrations with the same number of cells added in a reaction. When adding 500,000 HEK-293 cells to a 10 µL reaction, RT and PCR was inhibited by only ∼2 cycles at each step (**Fig. 6d**), comparable to the degree of inhibition with 6,000 Jurkat T cells. These mild losses were in quantitative agreement with the complete absence of an inhibitory regime we observed with *GAPDH* probes (**Fig. 6e**), suggesting that HEK cell lysate is ∼100-fold less inhibitory and thereby may be more amenable to RT-PCR in smaller droplets compared to Jurkat T cells.

The vast difference between two cell types motivated us to survey additional lines including bone marrow lymphoblasts (K-562), hepatocellular carcinoma cells (Hep G2), and murine macrophages (RAW-264) (**Fig. S5**). We found that K-562 cells displayed only minimal inhibition of PCR (**Fig. 6f**), while RAW-264 cells were most inhibitory to PCR, with >50,000 cells per 10 µL reaction essentially precluding amplification (**Fig. 6g**). Except RAW-264, all other cell lines tested showed significantly lower inhibition than Jurkat T cells. When 500,000 K-562 cells were added, the reaction displayed <1 cycle loss for both PCR and RT-PCR suggesting almost no inhibition of RT; addition of 500,000 Hep G2 cells inhibited PCR by ∼2 cycles and RT by ∼2 additional cycles, similar to HEK-293 cells (**Fig. 6h**). This surprising variability between cell lines was consistent across lysate concentrations and most pronounced at high concentrations (**Fig. 6i**).

Common strategies to improve stability and activity of reverse transcriptase and Taq polymerase (i.e., adding additional Mg^2+^, EGTA, and dithiothreitol), had little or even inhibitory effects (**Fig. S6a-c**). In addition, while prior work reported that adding single-stranded DNA-binding proteins (SSBs) can mitigate RT-PCR inhibition in 1 nL volumes,^8^ we found no improvement under conditions that mimic picoliter volumes (**Fig. S6d**). Together, our results suggest cell lysate inhibition of PCR and RT is cell-type dependent, yet is generally low enough to remain compatible with single-cell RT-PCR in 10-100 picoliter droplets.

### Picoliter-volume droplet one-step RT-PCR from a single cell

Our measured RT-PCR efficiencies in the plate-based assay suggest that there is no critical droplet size below which RT-PCR should be fully inhibited. To test whether single-cell RT-PCR can proceed in picoliter-volume DEs under optimized conditions, we co-loaded Jurkat T cells, one of the most inhibitory cell lines, with TaqMan RT-PCR reagents and probes for *GAPDH* transcripts and quantified TaqMan fluorescence before RT, after RT, and after either 20 or 50 PCR cycles (**Fig. 7**). Droplets exhibited no background TaqMan (FAM) fluorescence prior to thermal cycling, remaining stable even after several hours in solution (**Fig. 7a**). Upon heat lysis, cell-containing droplets began to show low background fluorescence (**Fig. 7b**), consistent with the remnant lysate-induced probe cleavage. Following 20 cycles of PCR, microscopy images of droplets showed increased FAM intensities, but this signal remained indistinguishable from background fluorescence in FACS analysis (**Fig. 7c**) and significantly lower than the false positive signals observed without heat inactivation (**Fig. 3a,b**).

**Figure 7:**
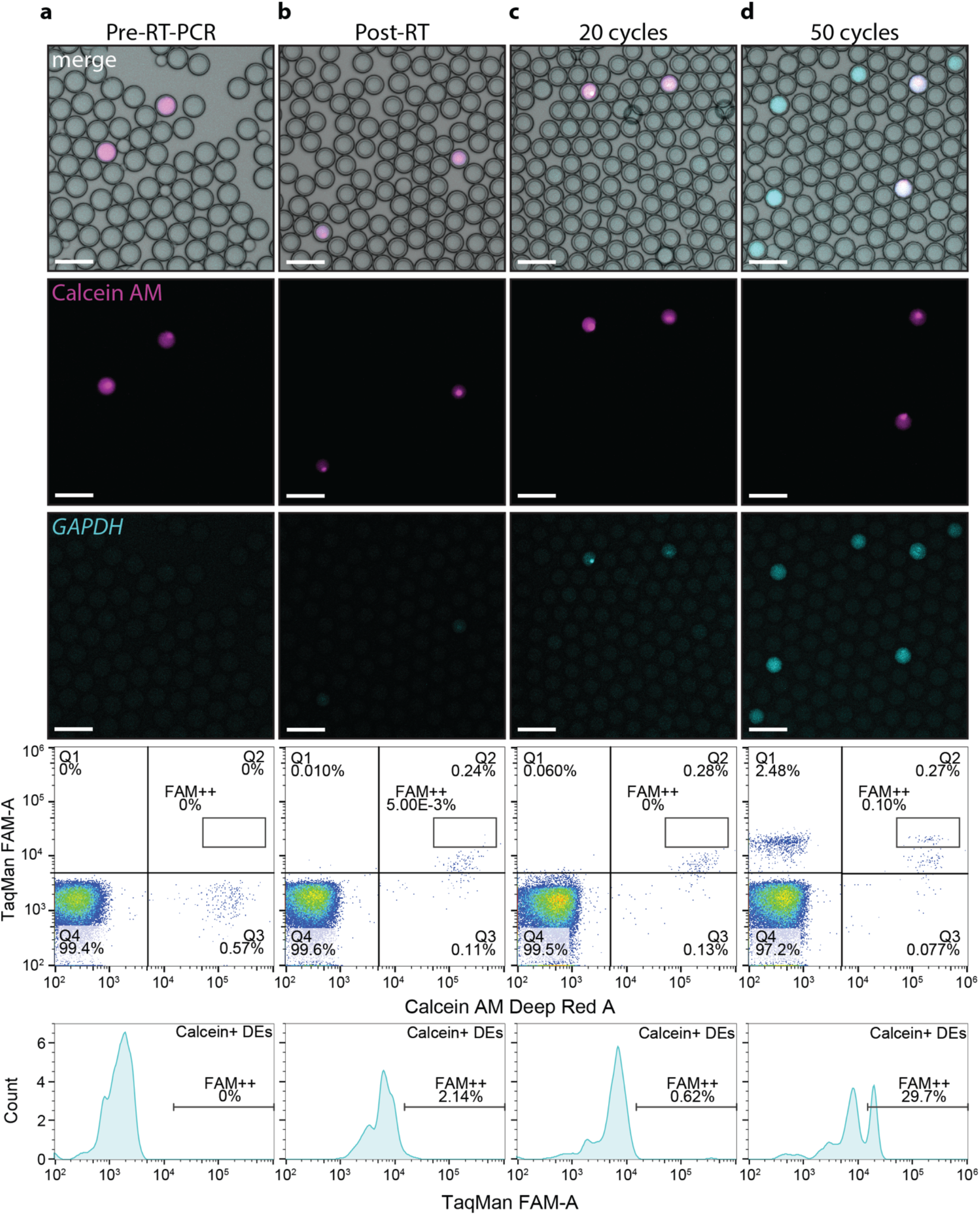
One-step RT-PCR from single cells encapsulated within 20 pL DE droplets. Representative bright field and fluorescence microscopy images (top: Calcein AM in magenta, FAM in cyan) and FACS plots (bottom) for DEs loaded with Calcein AM-stained Jurkat cells and TaqMan RT-PCR reagents for *GAPDH* after optimized RT-PCR (heat lysis, increased SuperScript III and Taq enzyme concentrations, and no lytic detergents or DTT). FACS density plots show TaqMan FAM signal *vs*. Calcein AM stain for DE-gated droplets; Q1, FAM^+^ Calcein^−^; Q2, FAM^+^ Calcein^+^; Q3, FAM^−^ Calcein^+^; Q4, FAM^−^ Calcein^−^ DEs. Histograms show distributions of FAM fluorescence in Calcein^+^ DEs, with the fraction of high FAM^++^ population denoted. Each column shows representative images and FACS plots at a different stage in the RT-PCR reactions: **(a)** DEs prior to RT-PCR, **(b)** DEs after heat lysis and RT but prior to PCR, **(c)** DEs after 20 PCR cycles, and **(d)** DEs after 50 PCR cycles. Microscopy images and FACS data reveal a population of cell-containing droplets that exhibit FAM fluorescence signal only after RT-PCR, indicating successful amplification of transcripts from single cells. Scale bars: 100 µm.

After 50 cycles of PCR, ∼30% of cell-containing droplets exhibited a distinct *GAPDH*^++^ population above that of background fluorescence that was easily visible both via microscopy and via FACS (**Fig. 7d**). As 50 cycles of PCR is an over-amplification—performed here to find the detection limit of true signal—we also observe FAM-positive, Calcein AM-negative droplets, signifying digital droplet PCR (ddPCR) amplification of individual stray transcripts released from loaded dead cells;^13^ however, these have a very low abundance, ∼2.5%, within droplets that did not contain a cell (**Fig. 7d**). Omitting both Taq and SuperScript III enzymes eliminated this *GAPDH*^++^ population, and background fluorescence levels did not change between the RT heat lysis step and amplification (**Fig. S7**). Together, these data represent, to our knowledge, the first successful demonstration of RT-PCR directly from a single cell in picoliter-volume reactions.

## Conclusions

The ability to sort and recover cells based on their RNA content could allow selective recovery of cell types of interest that cannot be isolated via FACS, including rare cell types revealed via single-cell transcriptomic studies^35^, cells containing latent integrated viruses,^36^ and cells expressing RNA splice isoforms or noncoding RNAs.^37^ While RT-PCR for single cells within encapsulated FACS-sortable double emulsion (DE) droplets provides a promising potential sorting method,^11-18^ realizing this technology requires successful RT-PCR from single cells within small volumes (<100 pL), at odds with many prior publications reporting that single-cell RT-PCR cannot take place at volumes <1 nL.^4-10^ Here, we used a plate-based assay designed to mimic conditions within DE droplets to optimize RT-PCR reaction conditions and enable amplification from single cells encapsulated within FACS-sortable DE droplets. The developed protocol should be broadly applicable to other picoscale reactors.^38^

Ultimately, downstream recovery of detailed transcriptomic information from single cells isolated based on RT-PCR signals will require the ability to integrate single-cell RNA-seq (scRNA-seq) library preparation into this pipeline. Although the temperature cycling required for TaqMan PCR may degrade labile RNA, inclusion of poly-dT primers during the initial reverse transcription step could convert all polyadenylated mRNA to the more stable cDNA prior to cycling. After FACS-based screening and isolation of fluorescent DE droplets within multiwell plates, scRNA-seq library preparation could proceed from recovered cDNA using standard techniques that employ well-based barcoding to group reads from individual cells. Nevertheless, our results suggest that RT and PCR steps may be sufficiently efficient for the preparation of single-cell RNA-seq libraries within sub-nanoliter double emulsion droplets, although additional studies will be required to determine detection sensitivity as a function of cell types, transcript abundance, and droplet size.

## Acknowledgements

We thank R. N. Hall for technical assistance. This work is supported by a HFSP grant RGY0085/2019 (B.W.) and NIH grant 1DP2GM123641 (P.M.F.). M.K. is a BioX SIGF Lavidge and McKinley Fellow and a Siebel Scholar. C.J.M. is a Damon Runyon Fellow supported by the Damon Runyon Cancer Research Foundation (DRG-2421-21). P.M.F. is a Chan Zuckerberg Biohub Investigator. B.W. is a Beckman Young Investigator.

## Supplementary figures

**Figure S1:**
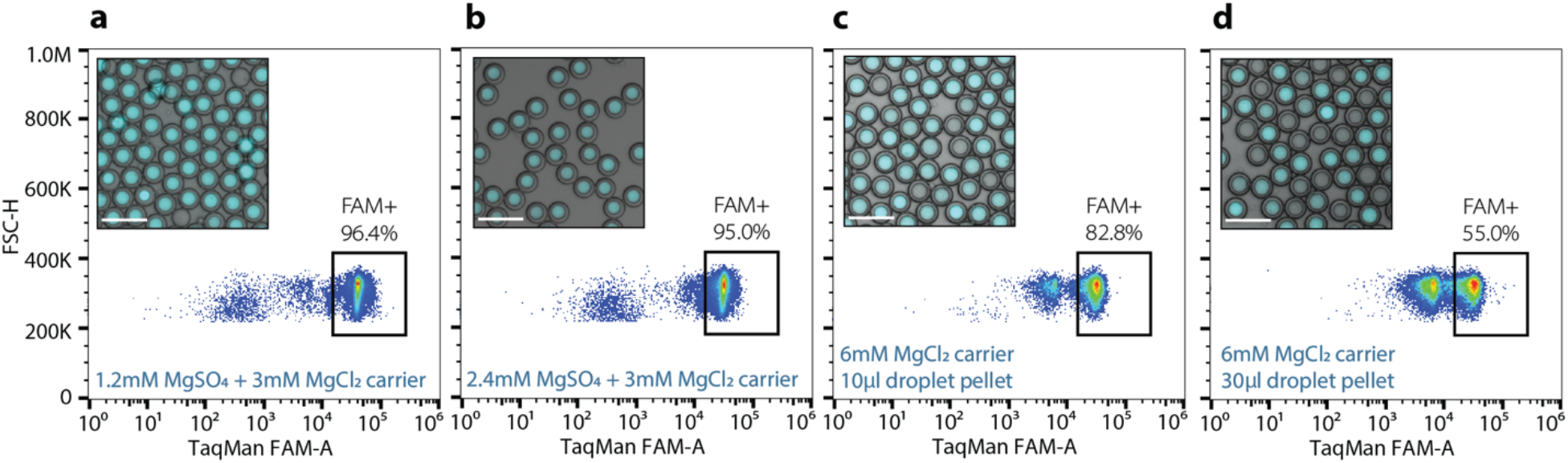
DE droplet RT-PCR requires additional Mg^2+^ salt in the carrier solution and needs to be performed in small volumes. Representative microscopy images (bright field and FAM fluorescence, merged) and FACS density plots for droplets loaded with 1 pg/droplet K-562 cell total RNA and RT-PCR reagents for detecting *GAPDH* transcripts after RT and PCR cycles when suspended in different outer aqueous carrier buffers: **(a)** 1× M-MLV buffer supplemented with 2.4 mM MgSO_4_, and **(b)** 1× Invitrogen One-Step RT-PCR buffer supplemented with 3 mM MgCl_2_. **(c, d)** Larger reaction volumes lead to smaller fractions of FAM^+^ droplets, potentially due to reduced Mg^2+^ accessibility from the carrier solution due to close packing of droplets. Scale bars: 100 µm.

**Figure S2:**
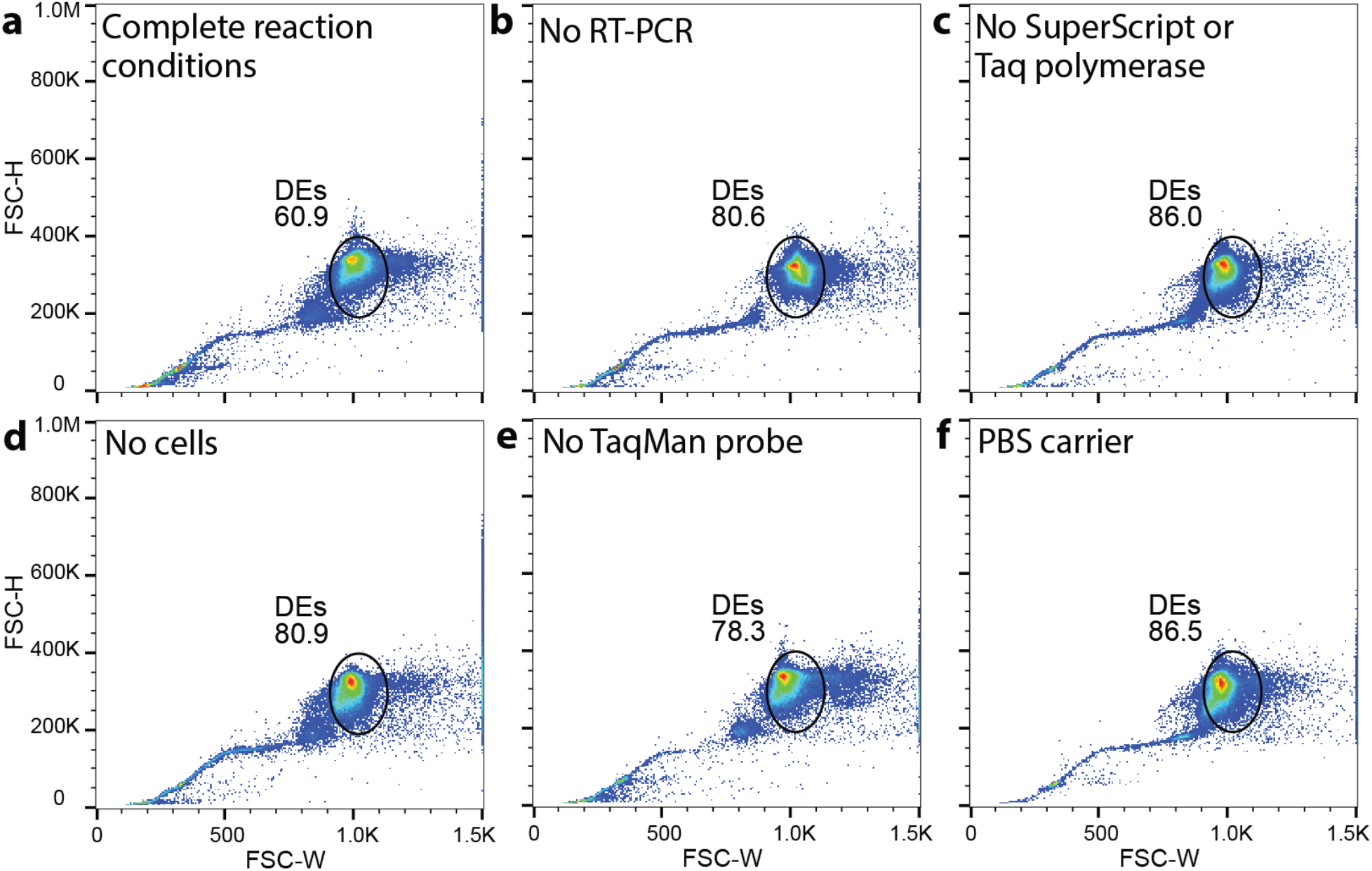
DE gates used in FACS analysis. FACS density plots show DE gates (black circles) and percentage of events within these gates for multiple reaction conditions, including: **(a)** Calcein AM Deep Red-stained Jurkat T cells and complete reaction buffer (*GAPDH* TaqMan RT-PCR reagents and lytic detergents: 0.1% NP-40, 0.1% Tween-20, 0.01% Digitonin) after RT\ and PCR thermal cycling, **(b)** Calcein-stained Jurkat T cells and complete reaction buffer without RT and PCR thermal cycling, **(c)** Calcein-stained Jurkat cells and reaction buffer without SuperScript reverse transcriptase and Taq polymerase, **(d)** complete reaction buffer alone without cells, **(e)** Calcein-stained Jurkat T cells and reaction buffer without TaqMan probe and primers, and **(f)** complete reaction buffer in a PBS outer carrier solution lacking Mg^2+^

**Figure S3:**
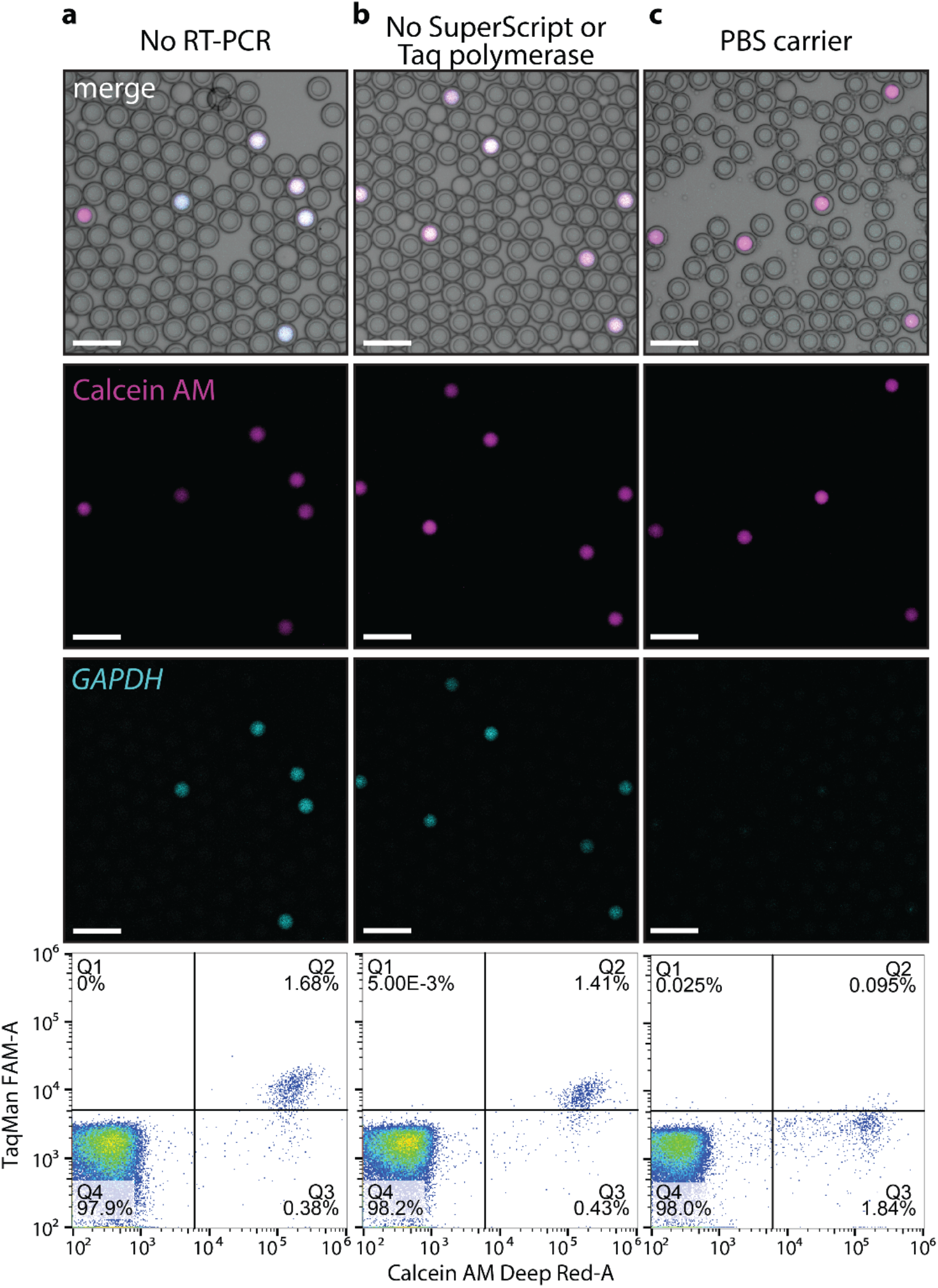
False positives of RT-PCR in DE droplets is still present without RT and PCR steps but absent when using carrier buffer without magnesium. **(a)** Calcein-stained Jurkat T cells and complete reaction buffer without RT and PCR thermal cycling, **(b)** Calcein-stained Jurkat T cells and reaction buffer without SuperScript reverse transcriptase and Taq polymerase, and **(c)** complete reaction buffer in a PBS outer carrier solution lacking Mg^2+^. FACS density plots display DE-gated droplets: Q1, FAM^+^ Calcein^−^; Q2, FAM^+^ Calcein^+^; Q3, FAM^−^ Calcein^+^; and Q4, FAM^−^ Calcein^−^. Scale bars: 100 µm.

**Figure S4:**
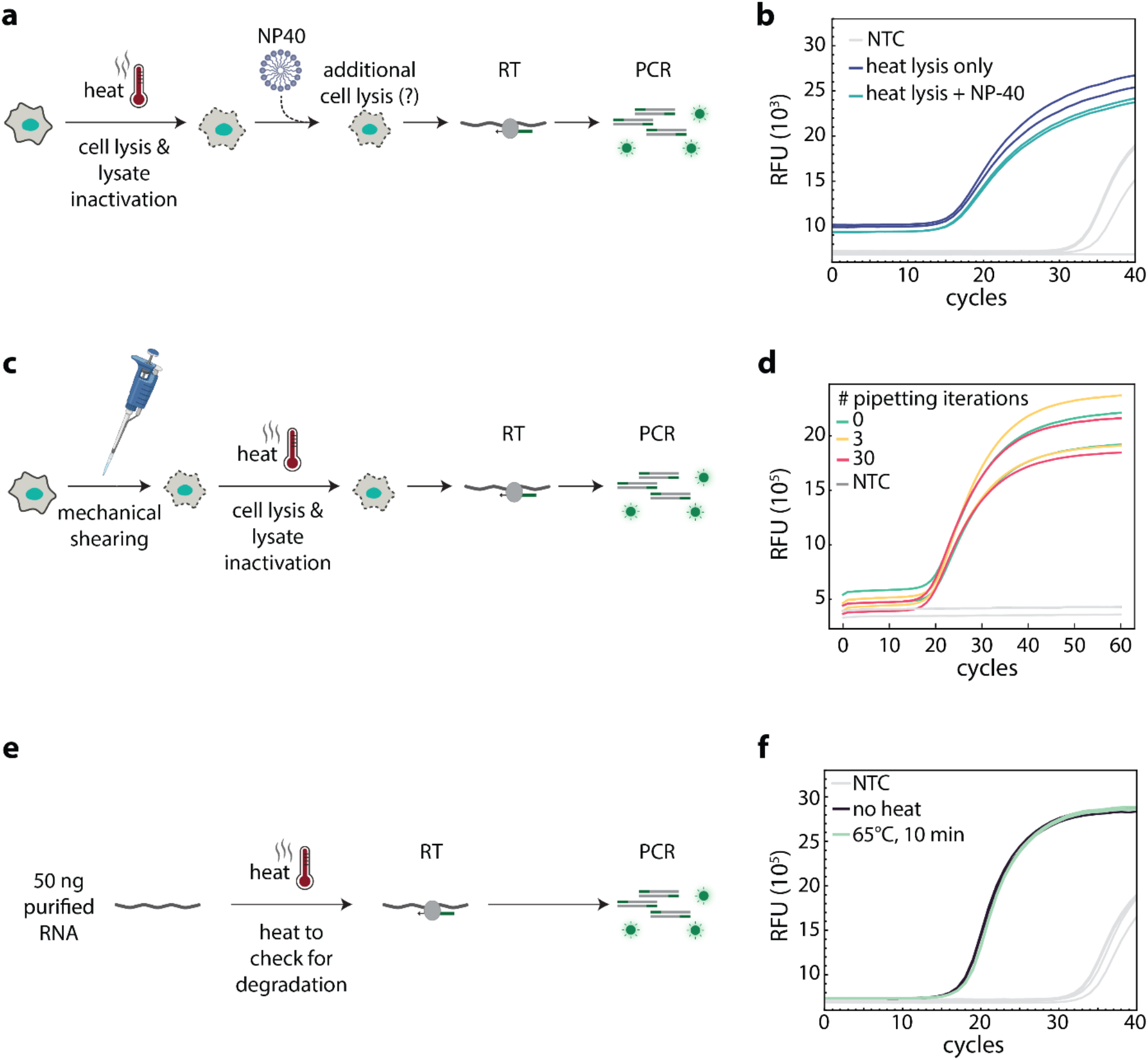
Reduced efficiency of RT-PCR is not due to insufficient lysis or RNA hydrolysis. (**a**) Schematic illustrating addition of NP-40 to test whether cells are fully lysed by heat. (**b**) qPCR traces for 500,000 Jurkat T cells in a 10 µL reaction to detect *GAPDH* transcripts with (blue line) and without (teal line) additional lytic detergents (0.1% NP-40). **(c)** Schematic illustrating mechanical shearing of cells prior to heat lysis. (**d**) qPCR traces for 50,000 Jurkat T cells in a 10 µL reaction to detect *GAPDH* transcripts with and without mechanical shearing prior to heat lysis. **(e)** Schematic illustrating heating of purified RNA to mimic conditions experienced during heat lysis to check for degradation. (**f**) qPCR traces for 50 ng purified RNA with (light green) and without (dark green) heat treatment. NTC: no template control.

**Fig. S5:**
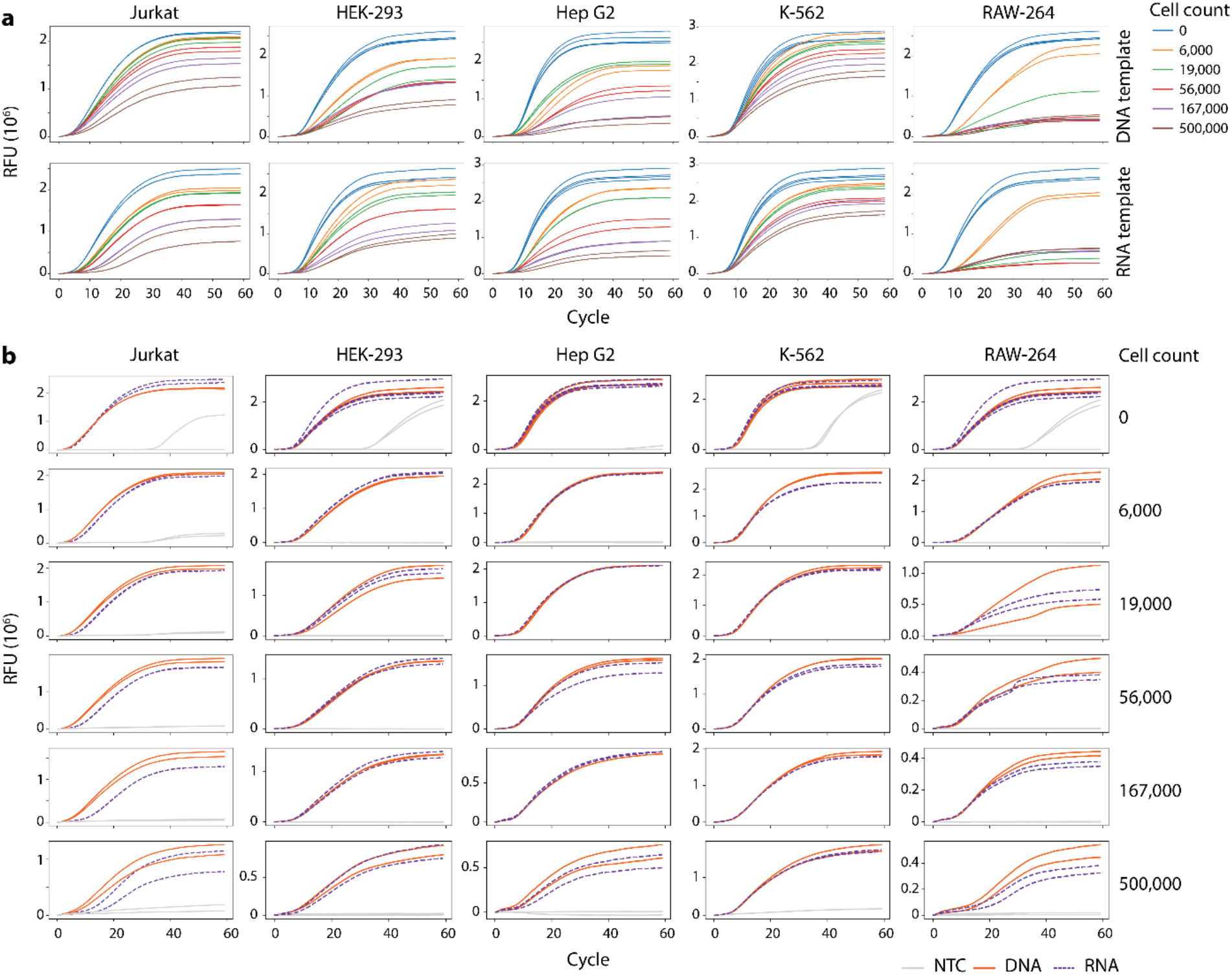
Lysate-induced inhibition of RT-PCR varies between cell lines. **(a)** Representative baseline-subtracted (RT)qPCR traces amplifying either DNA (top) or mRNA (bottom) *mTurquoise* templates in the presence of multiple added cell concentrations for five cell lines. **(b)** Data from (a), but with DNA (red) and mRNA (blue) curves overlaid for each cell line and cell concentration to directly compare of efficiency of PCR and RT-PCR. Grey lines are no-template controls (NTC).

**Figure S6:**
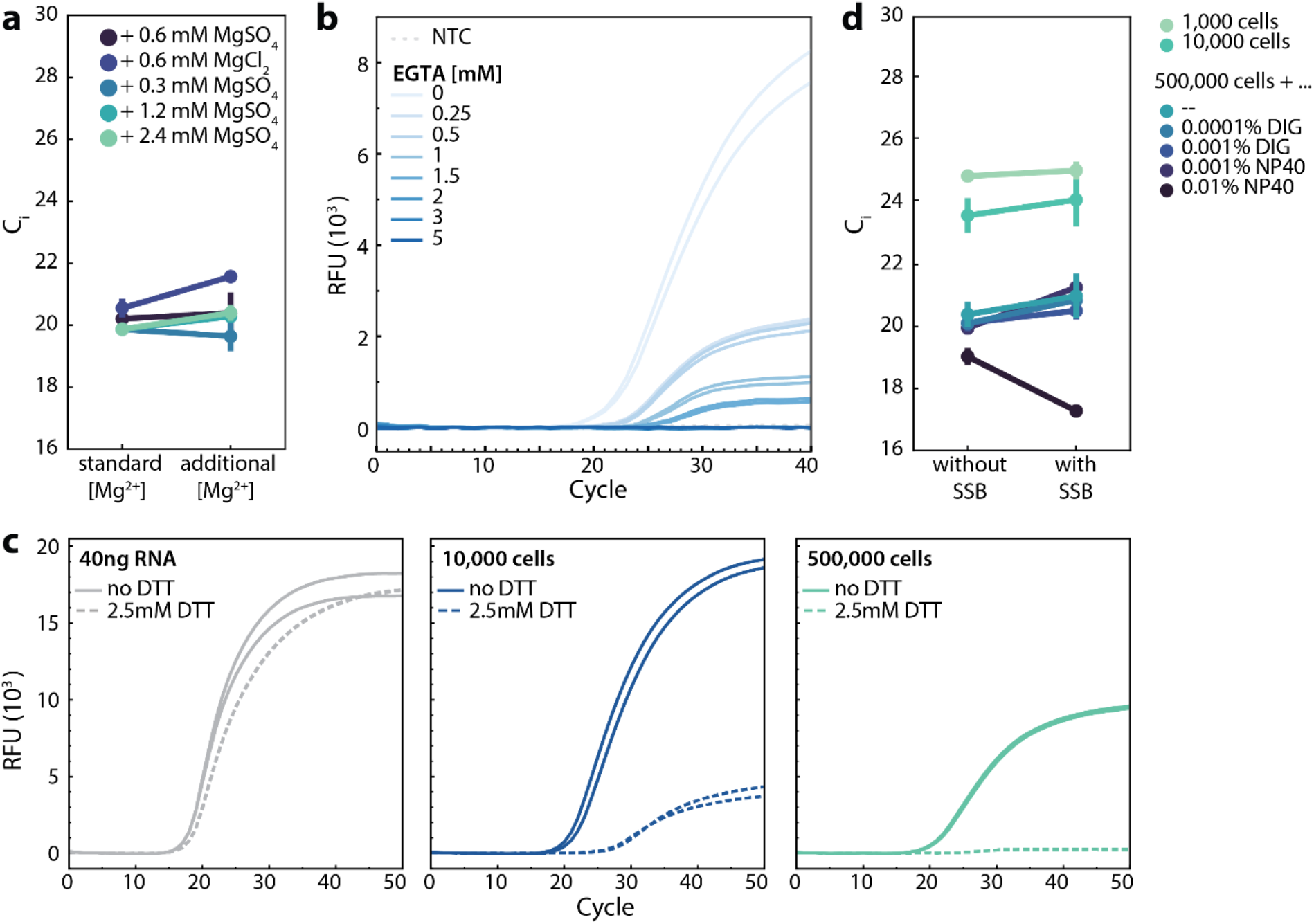
Cell lysate-induced inhibition is not improved with additional Mg^2+^, EGTA, DTT, or SSB protein. **(a)** The efficiency of RT-PCR reactions amplifying *GAPDH* transcripts (measured by C_i_) in the presence of different MgSO_4_ or MgCl_2_ concentrations with 500,000 Jurkat T cells in 10 µL reactions. The fluorescence signal dynamic range was not affected by adding Mg^2+^. Additional Mg^2+^ shows no significant benefit, indicating inhibition is not due to cofactor sequestration. **(b)** Baseline-subtracted qPCR traces amplifying *GAPDH* transcripts for 10 µL reactions with 500,000 Jurkat T cells in the presence of increasing concentrations of EGTA, which preferentially chelates calcium ions over magnesium and could reduce inhibition driven by competition of endogenous calcium for enzymatic Mg^2+^ cofactor binding sites. Added EGTA negatively affects amplification, indicating that endogenous calcium is not the source of reduced enzymatic efficiency. **(c)** Baseline-subtracted qPCR traces amplifying *GAPDH* transcripts for 10 µL reactions containing purified RNA, 10,000 cells, and 500,000 cells in the presence and absence of 2.5 mM DTT, which functions as a reducing agent to stabilize enzymes and is often added to RT and PCR reaction buffers. Added DTT has little effect on reactions with purified RNA but appears to inhibit RT-PCR with cells. **(d)** Efficiency of RT-PCR reactions amplifying *GAPDH* transcripts (measured by C_i_) for 1,000, 10,000, or 500,000 Jurkat T cells in 10 µL volumes in the presence and absence of lytic detergents NP-40 and Digitonin (DIG) and SSB protein. The fluorescence signal dynamic range was not affected by adding SSB.

**Figure S7:**
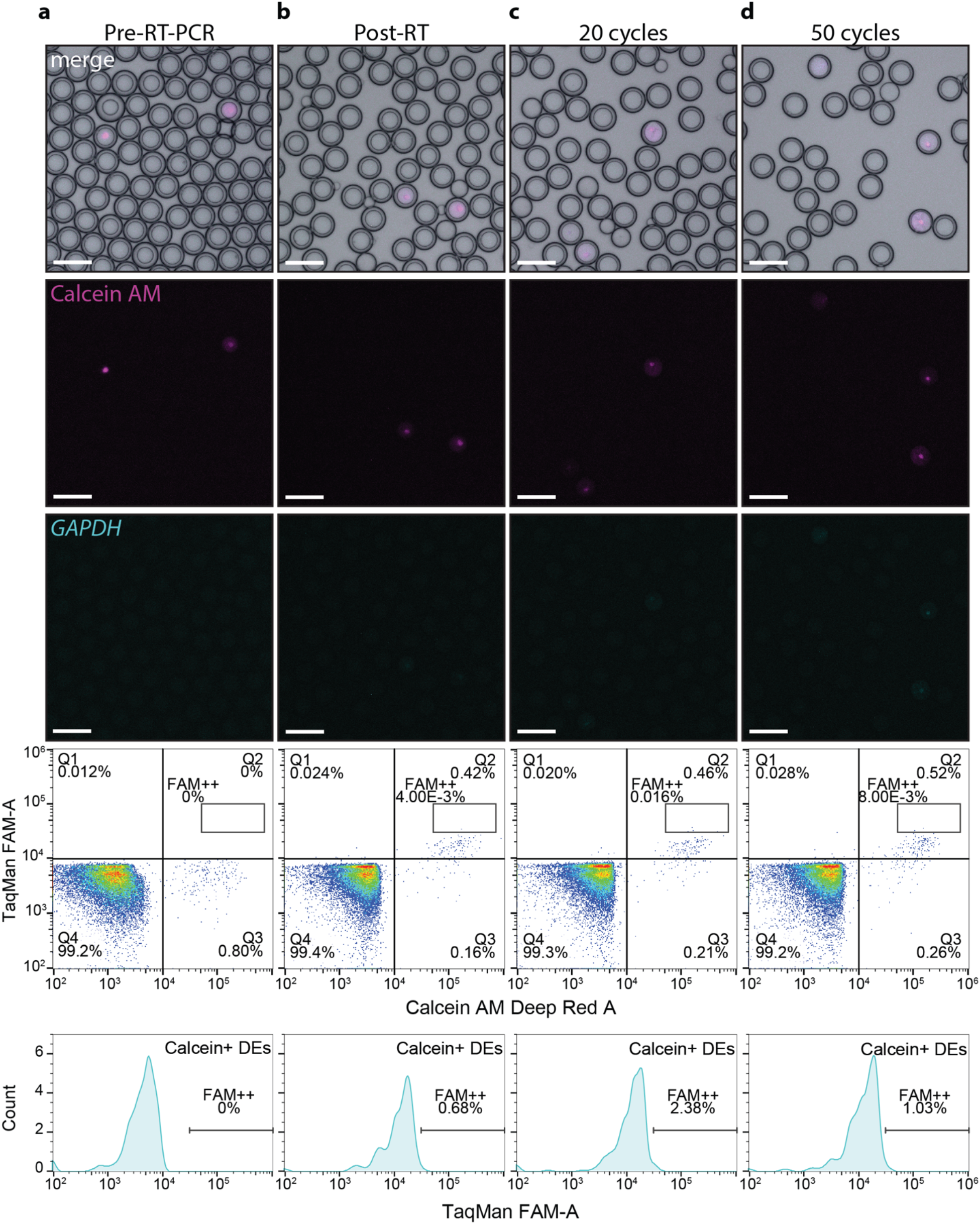
Optimized one-step RT-PCR protocol eliminates lysate-induced probe cleavage and false positives. Representative bright field and fluorescence microscopy images and FACS plots for DEs loaded with Calcein AM-stained Jurkat T cells and TaqMan RT-PCR reagents for *GAPDH* after RT-PCR in the absence of SuperScript III and Taq enzymes. FACS density plots show TaqMan FAM signal vs. Calcein cell stain for DE-gated droplets; histograms show distribution of FAM fluorescence in Calcein^+^ DEs, with the fraction of high FAM^++^ population denoted. Each column shows representative images and FACS plots at a different stage in the RT-PCR process: **(a)** DEs prior to RT-PCR, **(b)** DEs after heat lysis and RT but prior to PCR, **(c)** DEs after 20 PCR cycles, and **(d)** DEs after 50 PCR cycles. Unlike enzyme-containing reactions (**Fig. 7**), enzyme-free controls do not increase FAM signal above background after cycling. Scale bar: 100 µm.

